# EEG During Dynamic Facial Emotion Processing Reveals Neural Activity Patterns Associated with Autistic Traits in Children

**DOI:** 10.1101/2024.08.27.609816

**Authors:** Aron T. Hill, Talitha C. Ford, Neil W. Bailey, Jarrad A. G. Lum, Felicity J. Bigelow, Lindsay M. Oberman, Peter G. Enticott

**Affiliations:** Cognitive Neuroscience Unit, School of Psychology, Deakin University, Burwood, Australia; Centre for Human Psychopharmacology & Swinburne Neuroimaging, School of Health Sciences, Swinburne University of Technology, Melbourne, Australia; School of Medicine and Psychology, The Australian National University, Canberra, ACT, Australia; Monarch Research Institute Monarch Mental Health Group, Sydney, New South Wales, Australia; Noninvasive Neuromodulation Unit, Experimental Therapeutics and Pathophysiology Branch, National Institute of Mental Health, National Institutes of Health, Bethesda, MD, United States

**Keywords:** aperiodic activity, autism, connectivity, electroencephalography, oscillations

## Abstract

Altered brain connectivity and atypical neural oscillations have been observed in autism, yet their relationship with autistic traits in non-clinical populations remains underexplored. Here, we employ electroencephalography (EEG) to examine functional connectivity, oscillatory power, and broadband aperiodic activity during a dynamic facial emotion processing (FEP) task in 101 typically developing children aged 4-12 years. We investigate associations between these electrophysiological measures of brain dynamics and autistic traits as assessed by the Social Responsiveness Scale, 2nd Edition (SRS-2). Our results revealed that increased FEP-related connectivity across theta (4-7 Hz) and beta (13-30 Hz) frequencies correlated positively with higher SRS-2 scores, predominantly in right-lateralized (theta) and bilateral (beta) cortical networks. Additionally, a steeper 1/*f*-like aperiodic slope (spectral exponent) across fronto-central electrodes was associated with higher SRS-2 scores. Greater aperiodic-adjusted theta and alpha oscillatory power further correlated with both higher SRS-2 scores and steeper aperiodic slopes. These findings underscore important links between FEP-related brain dynamics and autistic traits in typically developing children. Future work could extend these findings to assess these EEG-derived markers as potential mechanisms underlying behavioural difficulties in autism.

## Introduction

The ability to accurately interpret complex emotional information from human faces is essential for successful social interaction, enabling an individual to infer the intentions and emotional states of others (Calvo & Nummenmaa, 2016; Collin et al., 2013). Facial emotion recognition is present within the first year of life and continues to develop throughout childhood, reaching maturity in adolescence or early adulthood (Leppänen & Nelson, 2008; Meinhardt-Injac et al., 2020; Walker-Andrews, 1998) coinciding with the extensive structural and functional changes that take place during these important neurodevelopmental periods (Aylward et al., 2005; Braams & Crone, 2017). Facial emotion processing (FEP) difficulties have been consistently associated with autism spectrum disorder (hereafter referred to as *autism*), a heterogenous neurodevelopmental condition that is characterised by impairments in social communication and interaction, restricted and repetitive behaviours, and atypical sensory processing (Lord et al., 2020; Uljarevic & Hamilton, 2013). Indeed, FEP impairments may be one of the earliest indicators of abnormal brain development in autism (Dawson et al., 2005), and research has repeatedly shown that the ability to interpret and understand others’ emotions is compromised in autistic individuals (Poljac et al., 2013; Uljarevic & Hamilton, 2013), with these difficulties present from early childhood (Fridenson-Hayo et al., 2016; Lacroix et al., 2014; Rump et al., 2009).

Beyond findings specific to autism, several studies have also identified links between FEP abilities and autistic traits in the broader, non-clinical population. This includes longitudinal research indicating that elevated autistic social difficulties in children are associated with poorer facial emotion recognition ability later in adulthood (Reed et al., 2021). Additionally, cross-sectional studies have shown that individuals with higher autistic traits exhibit delayed reaction times in tasks that require them to recognize others’ mental states (Miu et al., 2012) and display subtle difficulties in recognising emotional facial expressions in partially covered faces (Pazhoohi et al., 2021). Together, these findings broadly align with the clinical literature indicating impairments in FEP in autism (Fridenson-Hayo et al., 2016; Griffiths et al., 2019; Loth et al., 2018; Rump et al., 2009; Uljarevic & Hamilton, 2013). What remains less clear, however, are the precise neural mechanisms underlying disrupted FEP across non-clinical populations. This gap in understanding can be addressed by employing methods such as electroencephalography (EEG) to record brain activity during FEP tasks. EEG enables non-invasive recording of electrical neural activity produced by the brain using electrodes placed across the scalp and has high temporal resolution (millisecond timescale), and good tolerance to movement, making it well suited to studying developmental populations (Bell & Cuevas, 2012; Herve et al., 2022).

In autism, studies of altered neural activity during FEP have predominantly focussed on analysis of EEG-derived event-related potentials (ERPs), which represent brain responses both time- and phase-locked to a stimulus (Kappenman & Luck, 2012). The most consistent observations have been atypical modulation of the N170 potential, which is sensitive to faces and is theorised to reflect early encoding of facial stimuli (Bentin et al., 1996; Taylor et al., 2004). Specifically, reduced N170 amplitudes and/or delayed N170 latencies have been reported in several studies of autistic children, relative to neurotypical children (Batty et al., 2011; de Jong et al., 2008; Tye et al., 2014), with the N170 latency to upright human faces being accepted into the FDA’s Biomarker Quantification Program in 2019 (McPartland et al., 2020).

Presently, few studies have investigated EEG activity during FEP using methods other than ERPs. Amongst non-ERP studies, reduced frequency delta and theta power has been reported in autistic adults and adolescents (Tseng et al., 2015; Yang et al., 2011), while diminished theta connectivity in response to viewing emotional faces has been reported in autistic children, relative to neurotypical controls (Yeung et al., 2014). Additionally, using magnetoencephalography (MEG) data and taking a whole brain network-based approach, Safar et al. (Safar et al., 2018) found increased alpha-band connectivity in autistic children during processing of emotional (happy) faces; while reduced beta-band connectivity has also been reported in autistic children relative to controls during viewing of emotive faces (Safar et al., 2022). These limited findings suggest disrupted neural communication during FEP in autistic individuals. This observation aligns with more well-established reports of atypical functional connectivity patterns across brain networks in autism, which have been documented across multiple recording modalities including fMRI (Di Martino et al., 2013; Holiga et al., 2019; Ilioska et al., 2022), EEG (Murias et al., 2007; Shou et al., 2017; Zeng et al., 2017), and MEG (Lajiness-O’Neill et al., 2018; O’Reilly et al., 2017). However, the overall picture remains complex, with evidence for both hyper- and hypo-connectivity across various brain regions and networks (Ilioska et al., 2022; Uddin et al., 2013) (for recent reviews see: (Hull et al., 2016; Mehdizadefar et al., 2019; O’Reilly et al., 2017)).

In addition to capturing neural oscillations, the EEG signal also contains information on broadband aperiodic (i.e., non-oscillatory) activity (Donoghue et al., 2020; He, 2014). This arrhythmic scale-free signal demonstrates a 1/*f*-like spectral slope when examined in the frequency domain, whereby power diminishes exponentially with increasing frequency (Brake et al., 2024; He, 2014). Although initially regarded as ‘neural noise’, broadband aperiodic activity has received growing attention as a physiologically-relevant signal and potential marker of brain excitation-inhibition (EI) balance, given its sensitivity to drugs known to modulate excitatory (glutamatergic) and inhibitory (GABAergic) circuits (Colombo et al., 2019; Gao et al., 2017).

Specifically, a steeper aperiodic slope (i.e., larger exponent) has been associated with greater inhibitory tone (E<I), while a flatter slope suggests enhanced excitation (E>I) (Chini et al., 2022; Colombo et al., 2019; Gao et al., 2017; Waschke et al., 2021). We caution, however, that aperiodic activity is an emerging field of inquiry, with further research necessary to explore its relationship to complex EI systems in the brain (Brake et al., 2024). Nevertheless, given disrupted EI balance is currently a leading theory of autism (Rubenstein & Merzenich, 2003; Yizhar et al., 2011), characterizing differences in aperiodic slope is likely to be of significant value for establishing unique markers of atypical neural activity in this condition. To our knowledge, no EEG studies have investigated aperiodic activity in relation to autistic traits within non-clinical populations.

In this study, we used EEG recorded during a dynamic FEP paradigm (i.e., brief video clips of children’s faces) to assess associations between autistic traits and i) functional network connectivity, ii) aperiodic activity, and iii) neural oscillatory power in a cohort of typically developing children spanning early-to-middle childhood. These specific neurophysiological metrics were chosen given previous evidence of their disruption in autistic populations, as well as the putative link between aperiodic activity and EI balance (Chini et al., 2022; Gao et al., 2017; Murias et al., 2007; O’Reilly et al., 2017). We chose to examine dynamic, rather than static, stimuli as dynamic facial stimuli are potentially more sensitive and ecologically valid (Bernstein & Yovel, 2015; Zinchenko et al., 2018) producing robust activation of face-sensitive brain regions (Fox et al., 2009). Examining these neurophysiological metrics and their potential associations with autistic traits in a non-clinical sample also helps to avoid confounds related to co-occurring clinical diagnoses, which are common in autism (Bougeard et al., 2021; Lai et al., 2019), as well as any possible effects of psychotropic medications that can impact EEG recordings (Aiyer et al., 2016).

We hypothesised that autistic trait scores, as measured using the Social Responsiveness Scale, 2^nd^ Edition (SRS-2), would be associated with all three EEG-derived measures of neural activity – connectivity, aperiodic activity, and oscillatory power. However, due to the limited research in this area and the varied findings regarding EEG brain activity patterns during FEP, we refrained from making any specific directional predictions. Instead, we used data-driven approaches including the network-based statistic (NBS; connectivity) (Zalesky et al., 2010) to identify brain networks associated with autistic traits, and cluster-based permutation analyses (Maris & Oostenveld, 2007) to explore brain-wide links between oscillatory power or aperiodic activity and autistic traits.

## Materials and Methods

### Participants and Procedure

The data analysed in this study were collected as part of a larger project aimed at exploring cognitive function and electrophysiological activity across early-to-middle childhood (Bigelow et al., 2021, 2022; Hill et al., 2023; Hill et al., 2022a), but the electrophysiological data reported here (EEG recordings during a dynamic FEP task) have not been reported elsewhere. The initial sample included 153 typically developing children, as described by their primary caregiver, who were not diagnosed with any neurological or neurodevelopmental disorder. Out of the sample, 118 participants had complete SRS-2 assessments and task-related EEG recordings. The research received ethical approval from the Deakin University Human Research Ethics Committee (2017–065), while approval to approach public schools was granted by the Victorian Department of Education and Training (2017_003429). All EEG data were collected during a single experimental session, which was conducted either at the university laboratory, or in a quiet room at the participants’ school. SRS-2 caregiver reports were completed at each participating child’s home and then mailed to the investigators. Prior to commencement in the study, written consent was obtained from the primary caregiver of each child. Details of the experimental protocol were also explained to each child who then agreed to participate. Participant demographics are provided in Table 1.

**Table 1:** Participant demographics.

|  | Mean | SD | Range |
| --- | --- | --- | --- |
| Age (Years) | 9.78 | 1.71 | 4.1 - 12.9 |
| Sex (M:F) | 59:42 | - | - |
| SRS-2 T-Score | 48.88 | 8.98 | 37 - 85 |
| WASI FSIQ | 111.95 | 11.67 | 79 - 133 |

### Assessment of Autistic Traits

Autistic traits were evaluated using the School-Age version (ages 4–18 years) of the SRS-2, a 65-item caregiver report rating scale that measures deficits in social behaviour and restricted and repetitive behaviours associated with autism (Constantino & Gruber, 2005). The SRS-2 has strong psychometric properties and is one of the most widely used measures for characterising autism symptoms (Bruni, 2014). Results on the SRS-2 are reported as total severity scores, which were converted to *T*-Scores (Mean = 50, SD = 10), with higher scores indicative of more pronounced autism symptoms. The SRS-2 is comprised of a Total Score as well as five subscales reflecting more specific dimensions of autism-related symptoms (Social Awareness, Social Cognition, Social Communication, Social Motivation, and Restricted Interests and Repetitive Behaviour). For this study, we used Total *T*-Scores, rather than specific subscales, as these represent the most reliable measure of general social difficulties related to autism (Bruni, 2014).

### EEG acquisition and Facial Emotion Processing Task

EEG data were recorded via a 64-channel HydroCel Geodesic Sensor Net (Electrical Geodesics, Inc, USA) containing Ag/AgCl electrodes surrounded by electrolyte-wetted sponges. Recordings were taken in a dimly lit room using NetStation software (version 5.0) via a Net Amps 400 amplifier with a sampling rate of 1 KHz and an online reference at the vertex (Cz electrode). Electrode impedances were checked to ensure they were < 50 KOhms prior to recordings commencing (considered “low” impedance on Geodesic high-input impedance amplifiers). During EEG recordings, participants completed a FEP task (Figure 1) using stimuli taken from the Child Affective Facial Expression Stimulus Set (CAFE) (LoBue, 2014; LoBue & Thrasher, 2014). Following presentation of a fixation cross (500-750 ms), dynamic (1000ms) images of children’s faces expressing either happiness or anger were presented in a randomised order on a 55 cm computer monitor positioned 60 cm from the participant using the E-Prime software (Psychology Software Tools, Pittsburgh, PA). The stimuli began as neutral expressions before dynamically morphing into each emotional expression. At the end of each stimulus presentation, a blue box appeared around the image for 750 ms, signalling the participant to respond with a button press indicating that the subject was feeling either “good”, or “not good”. Timing of the blue box ensured that participant responses did not overlap with the stimulus presentation phase. During the study, participants were also presented with static visual stimuli, which were not analysed here (see (Bigelow et al., 2022) for event-related potential outcomes reported using static trials).

**Figure 1:**
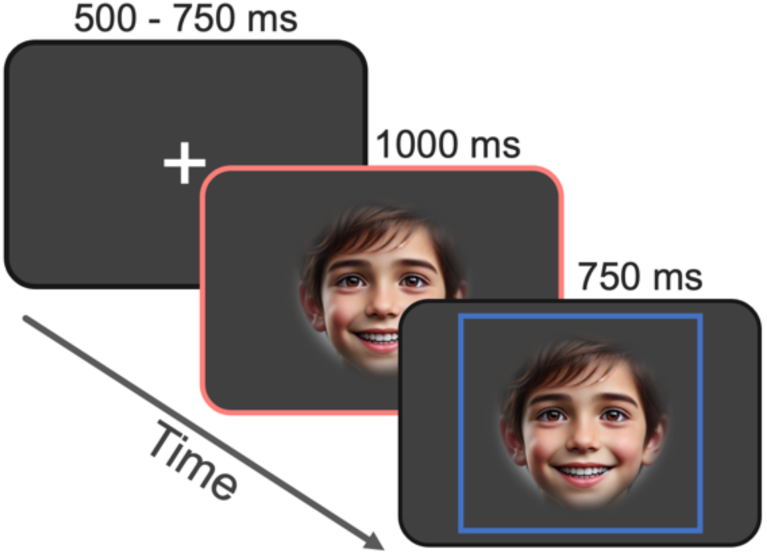
Example single trial of the dynamic facial emotion processing task. The middle image (highlighted in red) represents the dynamic stimulus presentation period, which was used for EEG analysis. Times (in milliseconds) denote the duration for which each stimulus appeared on the screen. Note: Face image is AI generated using DALL·E for this example.

### EEG Pre-Processing

The EEG data were pre-processed in MATLAB (R2021a; The Mathworks, Massachusetts, USA) using the EEGLAB toolbox (Delorme and Makeig, 2004) and custom scripts. The Reduction of Electroencephalographic Artifacts for Juvenile Recordings (RELAX-Jr) software (Hill et al., 2024) was used to clean each EEG file. This automated pre-processing pipeline was adapted from the RELAX software (Bailey et al., 2023) and is specifically optimised for cleaning data recorded in children using Geodesic SenorNet caps. It uses empirical approaches to identify and reduce artifacts within the data, including the use of both multi-channel Wiener filters and wavelet enhanced independent component analysis (ICA). The EEG data were first down-sampled to 500 Hz, then as part of the RELAX-Jr pipeline, data were bandpass filtered (0.25-80 Hz; fourth-order non-causal zero-phase Butterworth filter). Data were then notch filtered (47-53 Hz; fourth-order non-causal zero-phase Butterworth filter) to remove line noise, and any bad channels were removed using a multi-step process that incorporated the ‘findNoisyChannels’ function from the PREP pipeline (Bigdely-Shamlo et al., 2015) as well as multiple outlier detection methods from the original RELAX pipeline (including more than 20% of the electrode time series being affected by extreme absolute amplitudes, extreme amplitude shifts within each 1 second period, and improbable data distributions). Multi-channel Wiener filtering (Somers et al., 2018) was used to initially clean blinks, muscle activity, horizontal eye movement and drift, followed by robust average re-referencing (Bigdely-Shamlo et al., 2015). ICA was then computed using the Preconditioned ICA for Real Data (PICARD) algorithm (Ablin et al., 2018), with the adjusted-ADJUST IC classifier (Leach et al., 2020) used to select components for cleaning using wavelet-enhanced ICA (Castellanos & Makarov, 2006). Any electrodes rejected during the cleaning process were then interpolated back into the data using spherical interpolation (mean interpolated channels = 7.59, SD = 3.70). The cleaned data were further segmented from -1 to 1.5 seconds around the dynamic stimulus onset and any remaining noisy epochs (absolute voltages >120 µV, or kurtosis/improbable data with standard deviations [SD] >3 overall, or >5 at any electrode) were rejected (mean rejected epochs = 28.82, SD = 22.86 [11.68% of total dataset]). Finally, any participants with <50 trials were excluded (n = 17) to further ensure adequate signal-to-noise ratio and reliability for statistical analyses (Boudewyn et al., 2018; M.X. Cohen, 2014a). This left a total of N=101 participants with data included in the subsequent analyses (mean number of trials = 80.76, SD = 17.97). Participant demographics are provided in Table 1. Given the relatively wide age and IQ range across participants, we also checked for any possible associations between these values and SRS-2 *T*-scores using Pearson correlations. These revealed no significant associations between age and SRS-2 scores (r = -0.082, p = 0.415) or IQ and SRS-2 scores (r = -0.174, p = 0.084).

### Connectivity Analysis

Prior to performing the connectivity analyses the estimate of the scalp current density (surface Laplacian) was obtained from the EEG data using the spherical spline method, filtering out spatially broad features of the signal at each electrode, and thus better isolating neural activity under each electrode (Carvalhaes & de Barros, 2015; M. X. Cohen, 2014; Perrin et al., 1989; Srinivasan et al., 2007). This approach is recommended to protect against false positive inflation of connectivity measurement that can result from volume conduction (Miljevic et al., 2021). The EEG signal was then frequency transformed to obtain the complex Fourier coefficients for each subject/electrode across the 1 second window corresponding to the dynamic stimulus using a Hanning window, after which connectivity analysis was performed across the theta (4-7 Hz), alpha (7-13 Hz), and beta (13-30 Hz) bands using the weighted phase-lag index (wPLI) (Vinck et al., 2011).

The wPLI connectivity estimate, combined with the surface Laplacian spatial filter, assisted in reducing the prospect of confounds relating to volume conduction of the signal (Tenke & Kayser, 2015; Vinck et al., 2011). The wPLI method disregards instantaneous (i.e., zero phase-lag) interactions, which are characteristic of volume conduction, thus providing a more accurate connectivity estimate (Miljevic et al., 2021; Vinck et al., 2011).

### Calculation of the Aperiodic Signal and Oscillatory Power

To assess aperiodic and oscillatory activity, EEG data were first segmented into 1 second epochs corresponding to the duration of the dynamic stimulus. A Fourier transform (Hanning taper; 1 Hz resolution) was then applied across all channels for each participant to calculate spectral power. The spectral parameterization (*specparam*, formerly *fooof*) toolbox (version 1.0.0) (Donoghue et al., 2020) was then used to parameterise the Fourier transformed EEG data to extract the aperiodic spectral exponent (1/*f-*like broadband slope; frequency range: 1-40 Hz) independently for all electrodes. Models were fitted using the ‘fixed’ aperiodic mode, and spectral parameterisation settings for the algorithm were: peak width limits = [1, 8], maximum number of peaks = 12, peak threshold = 2, minimum peak height = 0.05. EEG power spectra were also calculated from the same 1 second epoch corresponding to the facial emotion stimulus. The aperiodic activity was then subtracted from the power spectra to leave only the oscillatory (periodic) component of the signal, which was then averaged across the theta (4-7 Hz), alpha (7–13 Hz), and beta (13– 30 Hz) bands ready for statistical analysis. This approach was taken to prevent conflating narrowband oscillatory activity with the broadband aperiodic signal (Brake et al., 2024; Donoghue et al., 2020; Donoghue et al., 2021). This was implemented using the Fieldtrip ‘ft_freqanalysis’ function within MATLAB, calling *specparam* functions from the Brainstorm toolbox (Tadel et al., 2011). We applied *specparam* using the same settings for the model as outlined for calculation of the aperiodic signal. Finally, since the calculation of power after spectral parameterisation to first remove the aperiodic signal is a relatively new approach, we also ran a traditional (i.e., non-parameterised) spectral analysis. This involved measuring the absolute EEG power spectra without subtracting the aperiodic activity (i.e., measuring the combination of periodic and aperiodic activity). Results from these analyses are reported in the Supplementary Materials (Figure S3).

### Statistical Anahlysis

Statistical analyses were performed in R (version 4.0.3; (R Core Team, 2020)) and MATLAB (version 2021a). Correlations between autistic traits (SRS-2 *T*-scores) and functional connectivity for each of the theta, alpha, and beta frequency bands were performed using the Network Based Statistic (NBS) MATLAB toolbox (Zalesky et al., 2010), which utilises non-parametric statistics in order to maintain statistical power while controlling for multiple comparisons (Zalesky et al., 2010). The primary threshold (test-statistic) for electrode pairs was set conservatively (test-statistic: 3.39, equivalent p-value of 0.001) to ensure that only robust connectivity differences would be compared at the cluster level and have strict control of Type 1 error (Wang et al., 2023; Zalesky et al., 2010). A value of p < 0.05 (two-tailed) was used as the secondary significance threshold for family-wise corrected cluster analysis (5000 permutations). Subsequent visualization of brain networks was performed using the BrainNet viewer toolbox (Xia et al., 2013).

Associations between SRS-2 *T*-scores and the aperiodic and periodic spectra were examined using non-parametric cluster-based permutation analyses in Fieldtrip using the ‘ft_freqstatistics’ function incorporating the ‘ft_statfun_correlationT’ function with SRS-2 score as the independent variable and the EEG data as the dependent variable (Maris & Oostenveld, 2007; Oostenveld et al., 2011). This approach allows for examination of global effects across all electrodes while controlling for multiple comparisons. For all comparisons, clusters were defined as > 3 neighbouring electrodes with a p-statistic < 0.05. Monte Carlo p-values (p< 0.05, two-tailed) were then subsequently calculated (5000 iterations). Lastly, we also performed experiment-wise multiple comparison controls using the Benjamini and Hochberg (Benjamini & Hochberg, 1995) false discovery rate (reported as *pFDR*) across the primary statistical tests comparing connectivity, aperiodic activity, and oscillatory power with SRS-2 scores (total of eight tests).

## Results

### Participant Demographics

The proportion of males to females did not differ significantly within the sample, *X*^2^ = 2.86, p = 0.097. Welch two-sample t-tests also confirmed that there was no difference between males and females in terms of age, *t*(93.00) = -0.39, *p* = 0.70, SRS-2 *T*-score, *t*(87.61) = -0.38, *p* = 0.71, or intellectual function as measured using the Wechsler Abbreviated Scale of Intelligence, Second Edition Full Scale IQ (WASI-FSIQ; conducted in participants aged ≥ 6 years), *t*(94.79) =− 0.63 *p* = 0.53. Density plots depicting the distribution of SRS-2 scores and age for males and females can be found in the Supplementary Material (Figure S1).

### Functional Connectihvity

NBS identified subnetworks that showed a significant correlation with autistic traits as measured using SRS-2 *T*-scores across both the theta (*pFDR* = 0.032) and beta (*pFDR* = 0.024) bands (Figure 2). No significant association was observed between alpha connectivity and autistic traits (*p* > 0.05). The theta subnetwork consisted of nine nodes (electrodes) and 12 edges (connections) spanning predominantly right parieto-temporal cortical regions. The beta subnetwork consisted of 15 nodes and 21 edges spanning bilateral (but predominantly right) frontal, temporal, and posterior regions. Further details, including all electrodes contributing to each subnetwork, are provided in the supplemental materials (Figure S2; Table S1). Next, we ran two additional multiple linear regression models with connectivity (wPLI; averaged across the electrode pairs forming the significant sub-network from NBS), age, and sex as predictors, and SRS-2 *T*-score as the outcome variable to further assess whether either age or sex could potentially also predict SRS-2 scores. For the theta band, the overall model was significant, *F*(3,97) = 11.84, *p* < 0.001, R² = 0.27. Of the predictors, only connectivity was found to significantly contribute to the model, *t*(97) = 5.86, *p* < 0.001. For the beta band, the overall model was significant, *F*(3,97) = 19.13, *p* < 0.001, R² = 0.37, with only connectivity found to significantly contribute to the model, *t*(97) = 7.49, p < 0.001.

**Figure 2:**
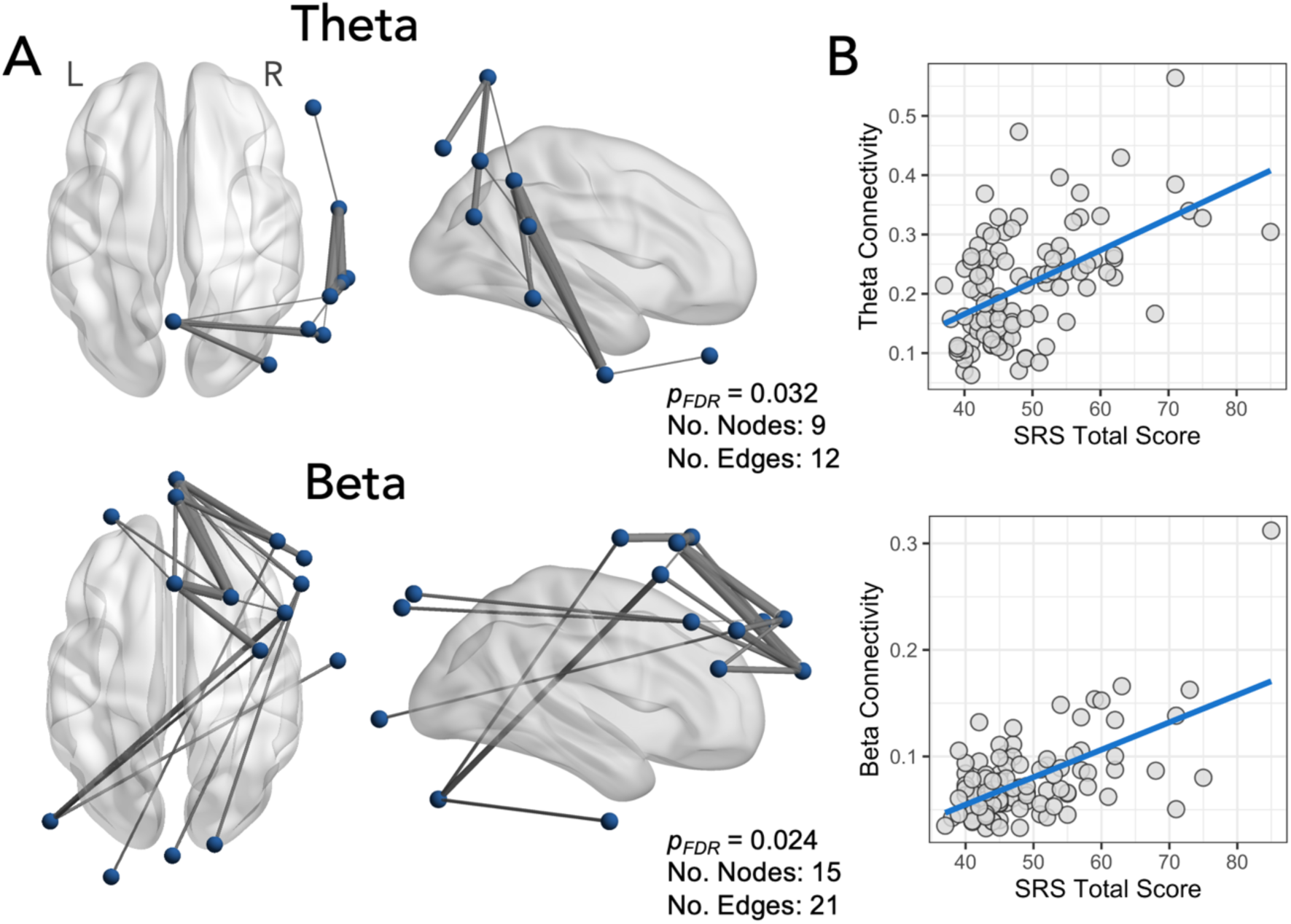
Associations between functional connectivity and level of autistic traits as measured by the Social Responsiveness Scale (SRS-2) total *T*-score. A) For the theta band, a significant connectivity subnetwork positively associated with SRS-2 score was identified spanning predominantly right temporo-parietal channels (top). For the beta band, a significant subnetwork positively associated with SRS-2 score was identified spanning both anterior and posterior channels (bottom). B) Accompanying scatterplots visually represent the association between connectivity strength (taken as an average of weighted phase-lag index values across the entire significant subnetwork) and SRS-2 scores.

### Aperiodic Slope

The spectral parameterisation algorithm performance was assessed using R^2^ and Error values, taken as an average across all electrodes, to determine the explained variance and error of the model fit relative to the power-frequency spectrum from each participant, respectively. Good model fits to the power-frequency spectrum were observed (mean R^2^ = 0.987, SD = 0.005; mean Error = 0.055, SD = 0.016), with all R^2^ values above 0.95 (Schaworonkow & Voytek, 2021). Cluster-based permutation analyses revealed a positive correlation between aperiodic slope and SRS *T*-scores (*pFDR* = 0.004) indicating steeper slopes in individuals with higher autistic traits. The positive cluster included electrodes spanning bilateral fronto-central regions (Figure 3; for specific electrodes forming the cluster see Supplementary Materials Table S1). No significant association was found between SRS-2 *T*-scores and aperiodic offset (*p* > 0.05, two-tailed). Next, as with the connectivity data, we ran additional linear regression models with aperiodic slope (taken as the average across all electrodes forming the significant cluster in the non-parametric permutation-based approach), age, and sex as predictors, and SRS-2 *T*-score as the outcome variable. The overall model was significant, *F*(3,97) = 6.49, *p* < 0.001, R² = 0.17. Of the predictors, only aperiodic slope was found to significantly contribute to the model, *t*(97) = 4.30, *p* < 0.001.

**Figure 3:**
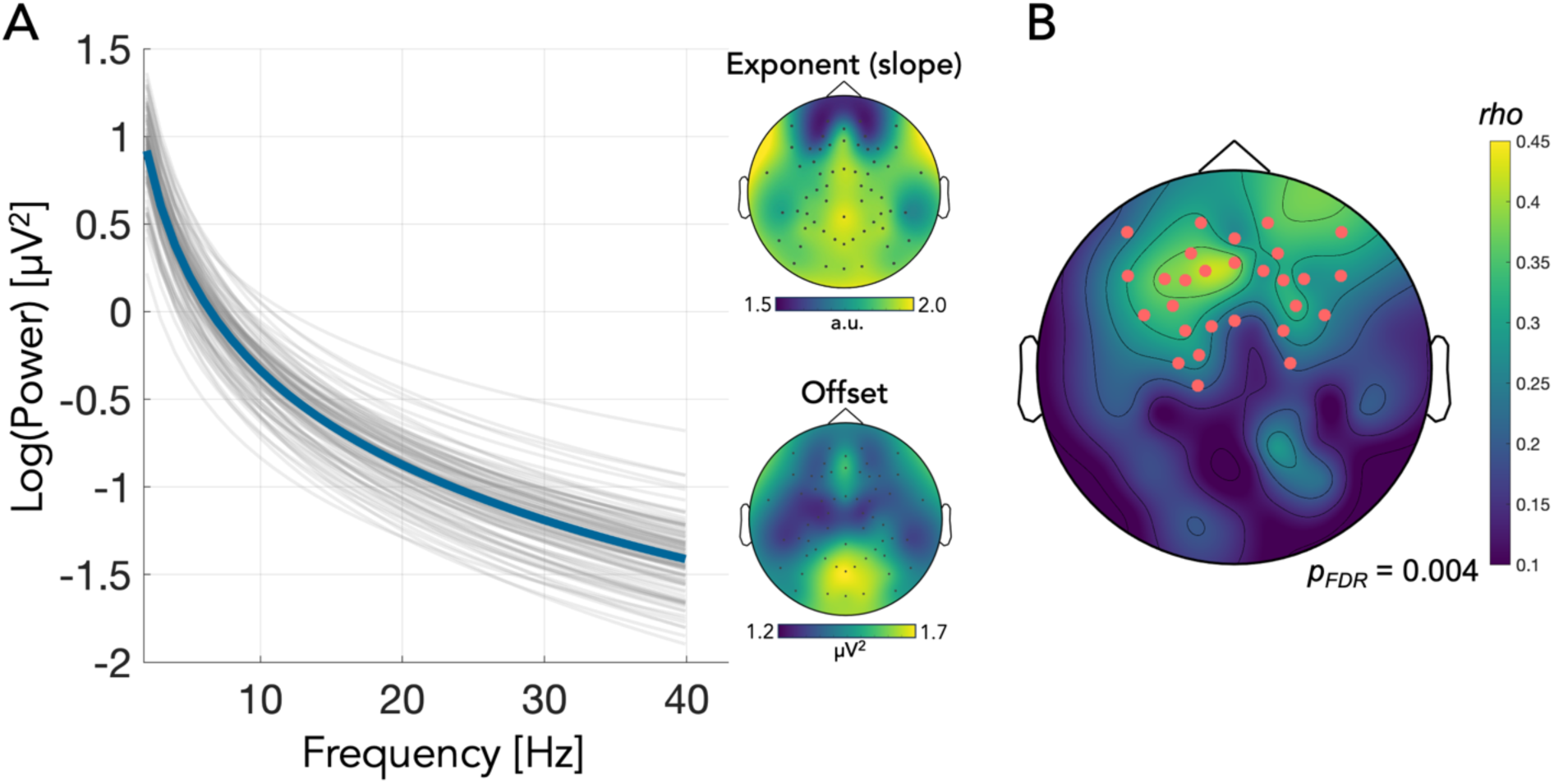
A) The aperiodic exponent (1/*f*-like slope). Thin grey lines represent the individual exponent values for each participant (taken as the average across all EEG channels), while the thick blue line represents the average exponent over all participants. The accompanying topographic maps show the distribution of exponent (top) and offset (bottom) values across the scalp. B) Topographic plot of the spatial distribution of the cluster of electrodes involved in the significant association between aperiodic slope and level of autistic traits as measured by the Social Responsiveness Scale (SRS-2) total *T*-score from the cluster-based permutation analysis.

### Oscillatory Power

Cluster-based permutation analyses were run to assess for associations between autistic traits (SRS-2 *T*-score; independent variable) and oscillatory power in each of the theta, alpha, and beta frequency bands (dependent variable). There was a significant positive association between SRS-2 score and power in the theta (*pFDR* = 0.024), and alpha (*pFDR* = 0.004) bands, but not the beta band (p > 0.05). The significant cluster within the theta band included predominantly fronto-parietal electrodes, while the alpha cluster incorporated electrodes with a broad scalp distribution, spanning bilateral anterior, central, and posterior regions (Figure 4; specific electrodes forming the clusters are provided in Supplementary Table S1.). As with the aperiodic slope, additional regression analyses were performed with oscillatory power (either theta or alpha; averaged across electrodes within the significant cluster), age, and sex as predictors, and SRS-2 *T*-score as the outcome variable. For the theta frequency, the overall model was significant, F(3,97) = 2.951, p = 0.036, R² = 0.08 with theta power being the only significant predictor contributing to the model, t(97) = 2.82 p = 0.006. For the alpha frequency, the overall model was also significant, F(3,97) = 6.20, p < 0.001, R² = 0.16, with alpha power being the only significant predictor contributing to the model, t(97) = 4.20 p < 0.001.

**Figure 4:**
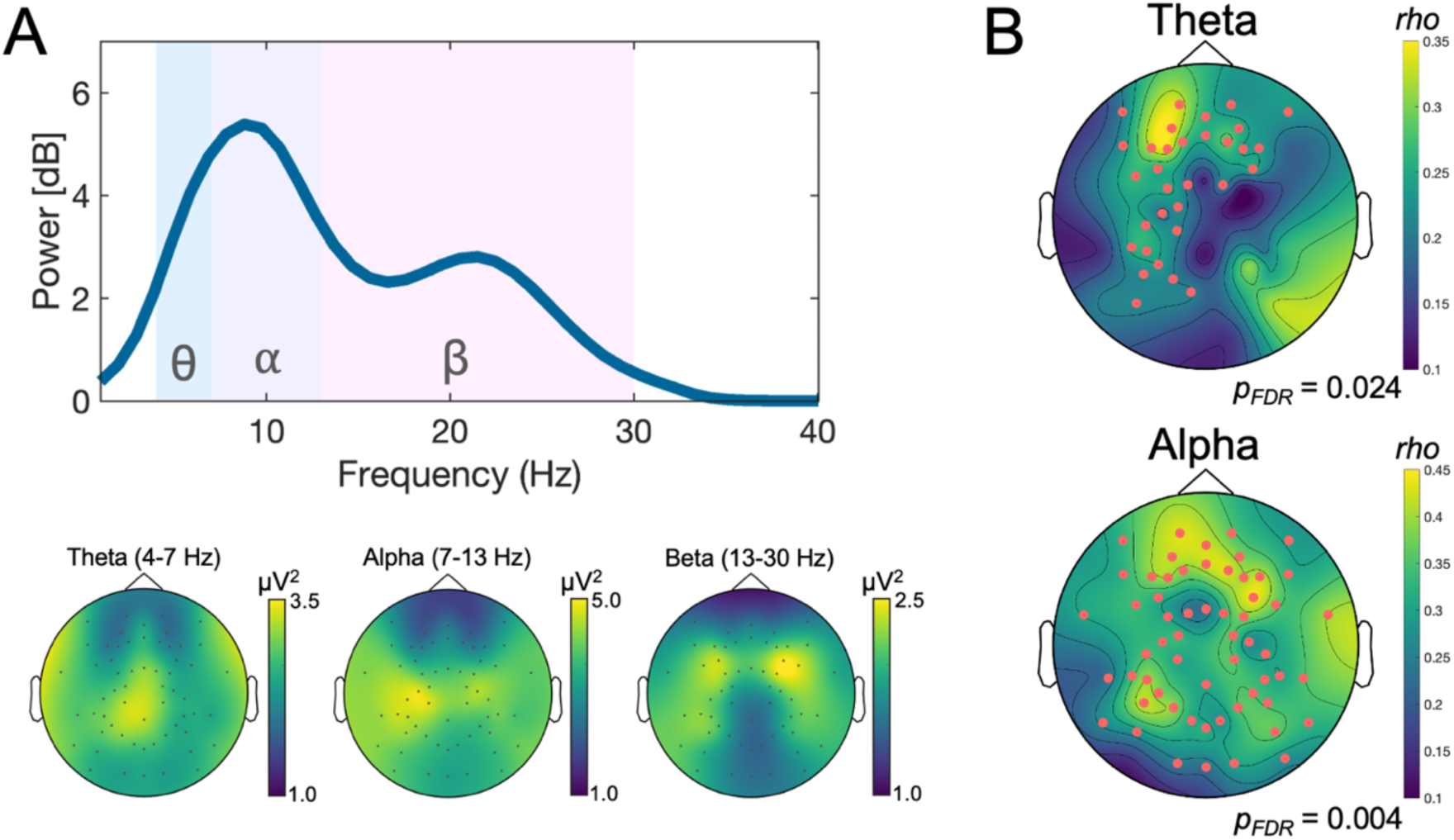
A) Periodic power spectra (i.e., after spectral parameterisation to remove the aperiodic signal) averaged across all electrodes (top) with topographic plots (bottom) showing the distribution of power across the scalp for the theta, alpha, and beta frequencies. B) Topographic plots of the spatial distribution of the electrodes forming the significant cluster for the association between theta (top) and alpha (bottom) power and Social Responsiveness Scale (SRS-2) *T*-score from the cluster-based permutation analysis.

Finally, cluster-based correlations were also performed using a more traditional approach taking spectral power without parameterisation (i.e., using data containing a combination of periodic and aperiodic activity). These results closely paralleled the findings from the parameterised periodic data (i.e., significant positive correlations with SRS2-2 *T*-scores), and are reported full in the supplemental materials (Figure S3).

As a final exploratory analysis, we also ran a correlation between theta and alpha power and aperiodic slope using the average signal across electrodes which comprised the significant clusters for each. The rationale for this was twofold. First, we wanted to test if our results could replicate similar recent findings from the literature reporting associations between these metrics (Hill et al., 2022a; Manyukhina et al., 2024; Merkin et al., 2023). Second, given alpha activity has been associated with inhibitory neuronal activity (Haegens et al., 2011; Mathewson et al., 2011) and has also been shown to be negatively correlated with BOLD fMRI activation (Gonçalves et al., 2006; Moosmann et al., 2003), we were interested in assessing if a relationship could be observed between power values and the aperiodic exponent, which has been linked to EI balance, with a steeper slope (larger exponent) potentially indicative of greater inhibition (E<I) (Gao et al., 2017; Waschke et al., 2021). Results indicated a significant moderate positive correlation (*rho* = 0.414, p < 0.001) between alpha power and aperiodic slope. There was also a weak but significant positive association between theta power and aperiodic slope (rho = 0.211, p = 0.034). Finally, as there was a single extreme outlier in the alpha power data (z-score > 3.29), we also re-ran the association after its removal; however, the association remained (rho = 0.412, p < 0.001). Correlation plots are provided in Figure 5.

**Figure 5:**
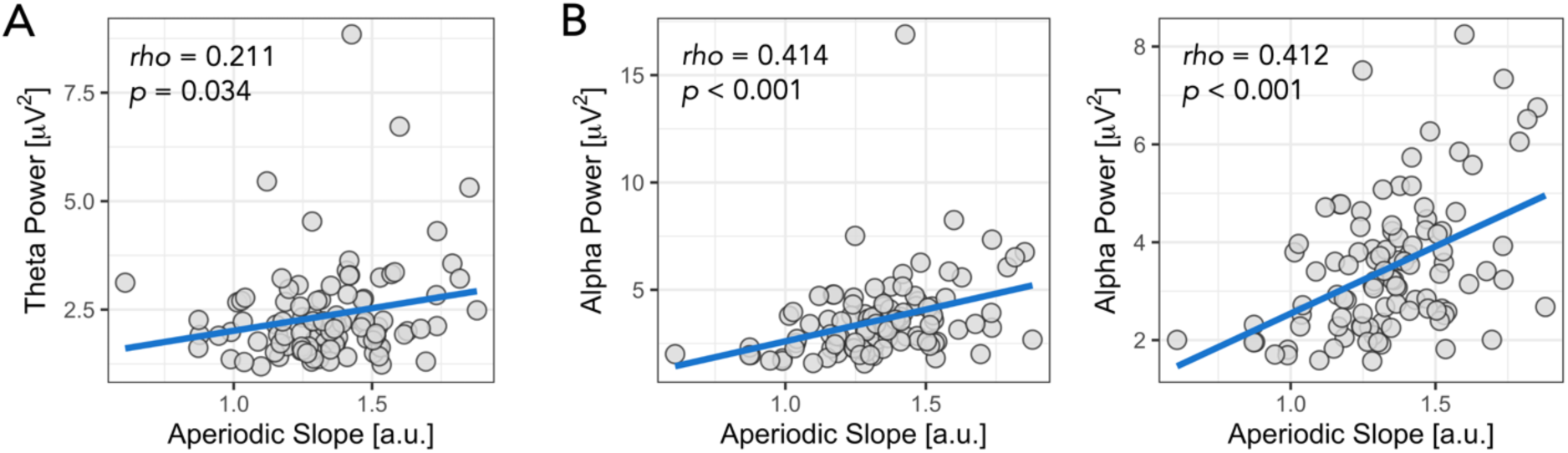
Association between aperiodic-adjusted theta (A) and alpha (B) power and the aperiodic slope. For alpha power, the plot on the left is the initial correlation, while the plot on the right is after removing the single extreme outlier.

## Discussion

Altered neural activity patterns, including atypical functional connectivity (Carson et al., 2014; Coben et al., 2008; Dickinson et al., 2018; Lajiness-O’Neill et al., 2018) and spectral power (Neo et al., 2023; Wang et al., 2013) have been repeatedly observed in autism. However, the association between neural activity and broader autistic traits in non-clinical populations remains under-investigated. Here, we sought to examine associations between autistic traits measured using the SRS-2 and EEG-derived measures of functional connectivity, spectral power, and aperiodic activity obtained from typically developing children while they were engaged in an ecologically valid social cognitive paradigm (a dynamic FEP task) (Arsalidou et al., 2011; Zinchenko et al., 2018). We found associations between SRS-2 total *T*-scores and EEG activity across all three metrics, highlighting important links between task-related brain activity patterns and autistic traits in typically developing children.

### Functional connectivity

We found a positive association between SRS-2 scores and functional connectivity within the theta and beta bands. Specifically, for each frequency, a subnetwork of stronger connectivity was identified to be associated with higher SRS-2 scores. In the theta band, this sub-network was lateralised to the right hemisphere, incorporating electrodes positioned over parietal, temporal, and frontal regions; while the beta sub-network was broader and bilaterally distributed, although predominately right-lateralised.

A dominant systems-level theory of autism, informed largely by resting-state fMRI research, suggests that autism is associated with atypical functional connectivity across brain circuits, including both hyper- and hypo-connectivity (Holiga et al., 2019; Ilioska et al., 2022). Interestingly, hyperconnectivity has been observed predominantly across network hubs located over prefrontal and parietal regions, corresponding closely to the central executive and default mode networks (DMN) (Buckner et al., 2008; Holiga et al., 2019; Ilioska et al., 2022). Hyperconnectivity between the DMN and other areas was also shown to be associated with social impairments (Ilioska et al., 2022). These observations appear to broadly align with our current findings, which identified subnetworks of stronger connectivity extending across electrodes over fronto-parietal regions, as well as broader central, temporal, and occipital areas in typically developing children with higher SRS-2 scores. Thus, our findings generally align with large-scale resting-state studies in the neuroimaging literature, which indicate disrupted connectivity across regions such as the DMN and fronto-parietal networks in individuals with autism (Holiga et al., 2019; Hull et al., 2016; Ilioska et al., 2022).

Nevertheless, it is important to acknowledge the limited spatial resolution of EEG (typically in the order of several centimetres), which restricts its capacity to offer precise insights into brain network activity (Ferree et al., 2001; Nunez et al., 1994). Future studies that integrate EEG with fMRI within the same sample could yield valuable complementary information regarding functional connectivity patterns across diverse temporal and spatial scales (Valdes-Sosa et al., 2011).

It is notable that the relationship between autistic traits and connectivity was predominantly observed in right hemispheric subnetworks, especially within the theta band. This finding is broadly consistent with prior fMRI studies, which have reported connectivity irregularities in these regions among individuals with autism (Hao et al., 2022; Igelström et al., 2016). These areas are also significantly involved in social cognition and face processing (Lombardo et al., 2011; Nomi & Uddin, 2015). Importantly, the present findings extend these observations beyond autistic populations to a sample of typically developing children showing a broad range of autistic traits. This suggests that the associations between autistic traits and connectivity extend across the broader autism phenotype, indicating that they are not exclusive to clinical populations. Further, while the majority of past literature has examined spontaneous resting-state network activity, here we provide insight into neural communication during a dynamic task that would be expected to strongly drive social-cognitive circuits within the brain (Fox et al., 2009). Moreover, the association between right-lateralised significant subnetworks and autistic traits supports hemispheric specialisation theories of emotion processing, consistent with neuroimaging findings showing greater right hemisphere activation during emotion processing (Ahern et al., 1991; Lindell, 2018; Sergent et al., 1992) (but see also: (Fusar-Poli et al., 2009)).

The literature on EEG/MEG connectivity patterns in autism is mixed, likely due to differences in experimental methodologies and participant heterogeneity. O’Reilly et al. (O’Reilly et al., 2017) systematically reviewed this topic, finding evidence for long-range hypoconnectivity in autism, especially in lower frequencies, with higher frequency bands showing mixed hypo- and hyperconnectivity. However, subsequent large-scale studies have failed to detect altered connectivity in autism (Garces et al., 2022). A resting-state EEG study by Aykan et al. (Aykan et al., 2021) found that right anterior theta connectivity could predict autistic traits (as measured by the Autism Spectrum Quotient [AQ]), with stronger connectivity associated with higher AQ scores; a finding we previously replicated using SRS-2 scores (Hill et al., 2022b). Here, we extend these findings by showing that both theta and beta connectivity during a FEP task are associated with SRS-2 scores.

### Aperiodic activity

We found that children with more pronounced autistic trait expression (i.e., higher SRS-2 scores) displayed steeper aperiodic slopes across fronto-central regions. Several recent pharmaco-EEG studies have demonstrated sensitivity of the aperiodic slope to drugs known to modulate either glutamatergic or GABAergic pathways, highlighting the aperiodic slope as a potential marker of EI balance within neural circuits (Colombo et al., 2019; Gao et al., 2017; Waschke et al., 2021). From an EI perspective, the present results might suggest greater inhibitory tone (E<I) during FEP in children exhibiting higher autistic traits.

Interestingly, this result appears to contrast with broader theories of EI imbalance in autism, which propose that increased excitation may be a possible underlying neurobiological mechanism (Rubenstein & Merzenich, 2003; Sohal & Rubenstein, 2019). One potential explanation for this difference is the variation in neural activity patterns between neurotypical individuals and the pathophysiological brain dynamics observed in those with a clinical diagnosis. Indeed, it has been recently shown that pharmacological challenge with the GABA agonist arbaclofen can produce divergent responses, in terms of aperiodic slope, between neurotypical and autistic individuals, whereby lower doses of arbaclofen can cause aperiodic slope to became steeper in autistic individuals, but elicit either a flatter slope, or no change, in neurotypical individuals (Ellis et al., 2023). It is also possible that alterations in EI systems are likely to show a degree of regional specificity. For example, autism animal models have shown both hyper- and hypo-excitability across various brain circuits (Antoine et al., 2019; Golden et al., 2018; Goncalves et al., 2017). Similarly, *in vivo* imaging of GABA and glutamate neuro-metabolites in humans using magnetic resonance spectroscopy (MRS) has also produced differing findings across several brain regions, showing both increases and decreases in glutamate (or Glx, a composite of glutamate + glutamine), and either decreases or no change in GABA (Ajram et al., 2019; Ford & Crewther, 2016). Future work using high-density EEG recordings or MEG combined with source-localisation approaches could be beneficial for examining aperiodic activity across specific brain regions (Hedrich et al., 2017).

Research into aperiodic activity in autistic populations is limited. Carter-Leno et al. found that steeper aperiodic slopes from EEG at 10 months correlated with higher autism traits (SRS-2 scores) at 36 months, but only in children with lower executive attention ability (Carter Leno et al., 2022). Using resting-state MEG recordings, Manyukina et al. (Manyukhina et al., 2022) reported flatter slopes in autistic boys (aged 6-15 years) with below-average IQ (IQ < 85) compared to neurotypical controls. However, no differences were observed in children with average IQs, and autistic children with an IQ above 85 tended to have steeper slopes, consistent with our observation of steeper slopes in individuals with higher autistic traits. Further research is needed to explore the mechanisms underlying these findings, however, one possibility, as suggested by Carter-Leno et al. (Carter Leno et al., 2022), is that steeper slopes may indicate a potential homeostatic compensatory mechanism, with enhanced inhibition in response to excessive excitatory activity to maintain stable EI ratios across neural circuits (Chen et al., 2022). Additionally, although aperiodic slope has been shown to reflect pharmacological modulation of EI systems (Colombo et al., 2019; Gao et al., 2017; Waschke et al., 2021), it likely also captures complex neural dynamics influenced by other physiological mechanisms (Brake et al., 2024). Future studies should therefore aim to better identify the primary neural generator(s) of the aperiodic signal.

### Neural Oscillations

In addition to aperiodic activity, we also found a positive association between oscillatory power in the theta and alpha bands and SRS-2 scores. The majority of EEG analyses examining oscillatory power have been performed using resting-state paradigms (Neo et al., 2023). Overall, this literature indicates a tendency for reduced alpha power in autism (Neo et al., 2023), with limited evidence for changes in theta frequencies. Although, results have shown considerable variability, possibly reflecting the large degree of heterogeneity present in autism, as well as variations in specific analysis methods used (Bogéa Ribeiro & da Silva Filho, 2023; Garces et al., 2022).

Task-related alpha oscillations likely reflect regulatory processes involved in inhibiting specific task-irrelevant brain regions (Herrmann et al., 2016; Jensen & Mazaheri, 2010). It is worth noting that instead of showing a region-specific relationship between alpha power during the task and autistic traits, our results indicated a broad topographical spread of greater alpha power related to higher SRS-2 scores. Therefore, one interpretation of our present findings is that the alpha-inhibitory mechanism is broadly heightened in individuals with higher autistic traits, perhaps reflecting widespread reductions in cortical processing in response to the dynamic facial stimuli. Somewhat contrastingly, however, Yang et al. (Yang et al., 2011) reported reduced occipital-parietal alpha power in a small sample of five adolescents and young adults with Asperger’s syndrome relative to controls during viewing of static emotive facial expressions (reported as greater event-related alpha desynchronisation using a time-frequency based approach) and interpreted this as a possible indicator of greater voluntary attention during face recognition. However, a later similar study, also in a small sample (n=10 Asperger’s, n=10 controls), reported no differences (Tseng et al., 2015). As our present results revealed enhanced alpha power in children with higher SRS-2 scores, it is possible that this might represent a tendency for reduced attention during the task in individuals with more pronounced autistic traits. Additionally, the association between theta power and SRS-2 scores was more confined to bilateral frontal and left parietal regions. Given the involvement of fronto-parietal networks and theta oscillations in top-down cognitive control (Cavanagh & Frank, 2014; Vuilleumier & Pourtois, 2007), we conjecture that this association might represent greater cognitive effort in processing the emotional stimuli in individuals with higher autistic traits. We also note that unlike previous studies, here we examined oscillatory power after first removing the aperiodic component of the signal, thus preventing the possibility of conflating periodic and aperiodic activity (Donoghue et al., 2020; Donoghue et al., 2021), but our results were maintained even when we did not control for the aperiodic activity.

Finally, we also observed positive associations between theta and alpha power and aperiodic slope values corroborating observations from several recent studies also showing relationships between aperiodic slope and periodic power (Hill et al., 2022a; Manyukhina et al., 2024; Merkin et al., 2023; Muthukumaraswamy & Liley, 2018). Given that both theta and alpha oscillations as well as steeper aperiodic slopes have been associated with inhibitory processes, we tentatively interpret this association as reflecting a shared underlying mechanism (i.e., cortical inhibition) contributing to these two EEG-derived metrics (Colgin, 2013; Gao et al., 2017; Jensen & Mazaheri, 2010; Manyukhina et al., 2024; Mathewson et al., 2011; Zhu et al., 2023). Considering the substantial value of dependable non-invasive markers for neural inhibitory processes in both cognitive and clinical neuroscience, it would be beneficial for future research to delve deeper into these associations and investigate how they respond to pharmacological and device-based interventions.

### Limitations

There were several limitations in the present study. First, as we combined emotional conditions across the FEP task, we cannot comment on effects as they might relate to the specific emotional valence of the stimuli. However, our primary objective was to investigate EEG activity and its association with autistic traits while individuals were engaged in FEP, rather than attempting to disentangle differences between specific emotions. Importantly, combining data across stimuli enabled us to ensure an adequate number of trials were included for each participant, thus helping to maximise data quality (i.e., adequate signal-to-noise ratio), and thus help minimise potential for spurious findings (M.X. Cohen, 2014b). Nevertheless, future studies could further investigate possible associations between autistic traits and specific emotional conditions. Second, we ran our analyses at the electrode level, rather than using a source-localised signal.

Thus, our results cannot provide fine-grained information on precise cortical or sub-cortical regions. As we did not collect structural MRI data for these participants nor individualised spatial coordinates of the electrodes, we chose not to run source estimation of the EEG signal to avoid the reduced reliability of this approach. Finally, we also note that while this was a typically developing population (i.e., participants were described as typically developing by their primary caregiver), it nevertheless still included some participants who obtained SRS-2 scores indicative of a degree of deficiency in social functioning. Given this, in addition to the developmental nature of the cohort, we therefore cannot rule-out the possibility that some participants might have later received a diagnosis of autism. However, having a broad range of SRS-2 scores is also a possible strength, as it captures a wider range of social differences.

## Conclusion

Using EEG activity recorded during a dynamic FEP task, we have demonstrated associations between functional connectivity, as well as periodic and aperiodic neural activity and autistic traits in typically developing children spanning early-to-middle childhood. These findings compliment past research examining functional brain processes in autism while extending these findings to a non-clinical sample. Collectively, these results highlight important connections between neural activity patterns and autism traits that extend to the broader population of typically developing children. These results also provide directions for future research to further elucidate the role of neurophysiological processes in autism, which includes the search for reliable biomarkers to gauge therapeutic outcomes and the detection of mechanisms that might further our understanding of autism pathophysiology.

### Funding

This work was supported by an Australian Research Council Future Fellowship (PGE; FT160100077). LMO is supported by the NIMH Intramural Research Program (ZIAMH002955).

