## Supplementary Material for "EEG During Dynamic Facial Emotion Processing Reveals Neural Activity Patterns Associated with Autistic Traits in Children"

### Supplementary Materials

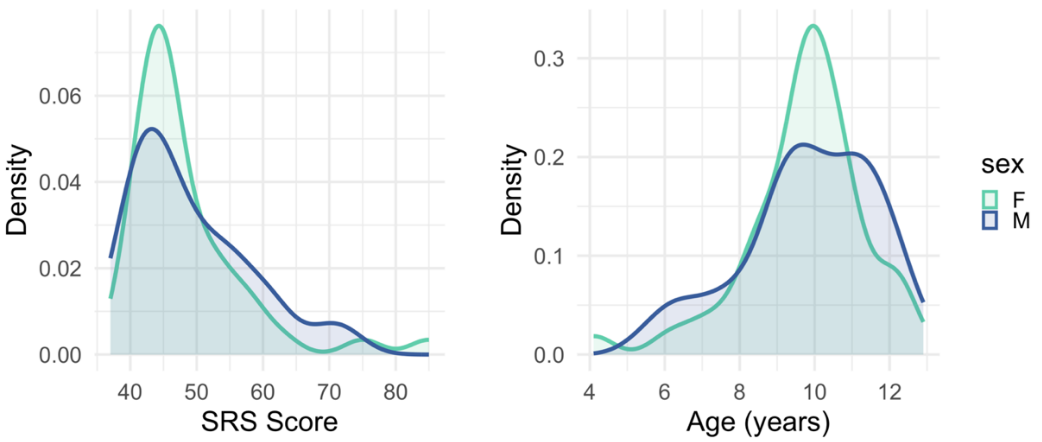


**Figure S1:** Density plot showing the distribution of SRS scores (total t-score; left) and age (right) for females and males. There were no significant differences in SRS score or age between the two sexes ($p > .05$).

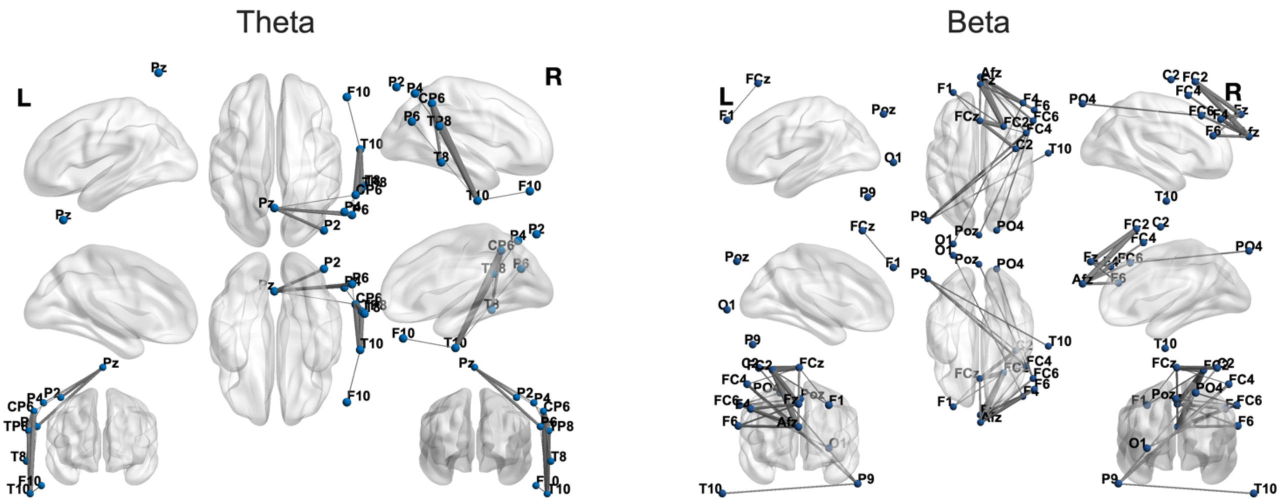


**Figure S2:** Plots of the significant subnetworks comparing functional connectivity (wPLI) and autistic traits (SRS-2 total *T*-scores) for the theta (left) and beta (right) frequency bands using the Network Based Statistic.

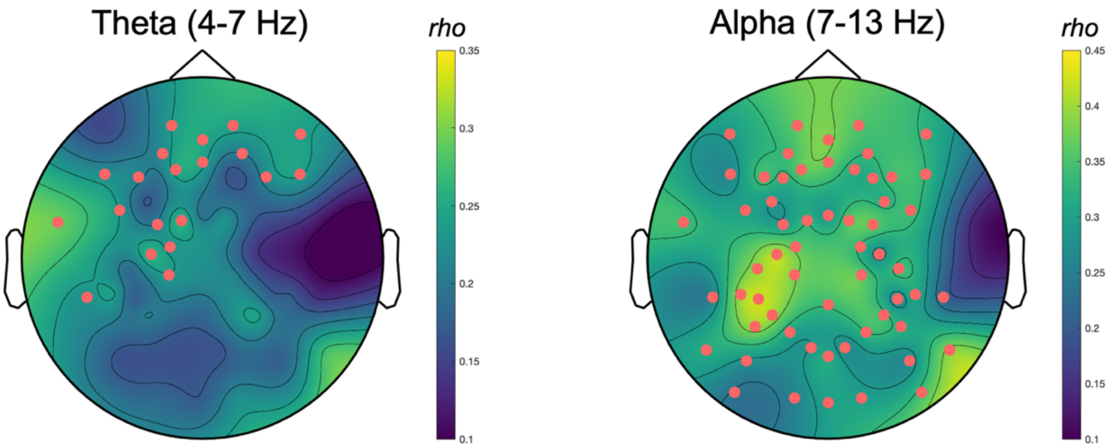


**Figure S3:** Topographic plots showing the significant cluster for correlations between total power (i.e., non-parameterised, including both aperiodic and periodic components of the signal) in the theta (4-7 Hz) and alpha (7-13 Hz) bands and SRS *T*-score. Similar to the findings reported in the manuscript, which used a spectral parameterisation approach to account for the aperiodic signal, there were significant positive correlations for both theta (p=0.022) the alpha band ($p < 0.001$) when using this traditional approach.

**Table S1:** Electrodes forming significant clusters (aperiodic activity, oscillation power) or networks (connectivity; electrode pairs) for correlations between EEG activity and SRS-2 *T*-scores.

| **Measure** | **Electrodes** |
|---|---|
| Aperiodic Slope | F10, AF4, F2, FCz, Fp2, Fz, FC1, AFz, F1, Fp1, AF3, F3, F5, FC5, FC3, C1, F9, F7, FT7, C3, CP1, C4, FC4, FT8, FC6, F8, F6, F4 |
| Theta Power | F10, AF4, F2, Fp2, Fz, FC1, AFz, F1, Fp1, AF3, F3, F5, FC5, FC3, C1, F9, F7, FT7, C3, CP1, C5, TP7, CP5, P5, P3, P7, P1, PO3, FC2, FC6, F6, F4 |
| Alpha Power | F10, AF4, F2, FCz, Fp2, Fz, FC1, AFz, F1, Fp1, AF3, F3, F5, FC5, FC3, C1, F9, F7, FT7, C3, CP1, C5, T9, T7, TP7, CP5, P5, P3, TP9, P7, P1, PO3, Pz, O1, POz, Oz, PO4, O2, P2, CP2, P4, P10, P8, P6, CP6, TP10, TP8, C6, C4, C2, T8, FC4, FC2, T10, FT8, FC6, F8, F6, F4 |
| Theta Connectivity | Pz-P2, Pz-P4, Pz-P6, Pz-CP6, P4-TP8, CP6-TP8, CP6-T8, TP8-T8, F10-T10, P6-T10, CP6-T10, TP8-T10 |
| Beta Connectivity | FCz-Afz, FCz-F1, FCz-C2, P9-C2, FCz-FC4, Afz-FC4, P9-FC4, FCz-FC2, Fz-FC2, Afz-FC2, F1-FC2, P9-T10, Fz-FC6, Afz-FC6, Poz-FC6, PO4-FC6, Fz-F6, Afz-F6, Fz-F4, Afz-F4, O1-F4 |
